# Competitive performance and superior noise robustness of a non-negative deep convolutional spiking network

**DOI:** 10.1101/2023.04.22.537923

**Authors:** David Rotermund, Alberto Garcia-Ortiz, Klaus R. Pawelzik

## Abstract

Networks of spiking neurons promise to combine energy efficiency with high performance. However, spiking models that match the performance of current state-of-the-art networks while requiring moderate computational resources are still lacking. Here we present an alternative framework to deep convolutional networks (CNNs), the ”Spike by Spike” network (SbS), together with an efficient backpropagation algorithm. SbS implements networks based on non-negative matrix factorisation (NNMF), but uses discrete events as signals instead of real values. On clean data, the performance of CNNs is matched by both NNMF-based networks and SbS. SbS are found to be most robust when the data is corrupted by noise, specially when this noise was not seen before.

## 1 Introduction

In this paper, we compare the performance of three different model classes for deep learning: standard deep convolutional neural networks (CNNs) [2], deep networks based on non-negative matrix factorization (NNMF) [5] [6], and a recent deep spiking neuronal network model called spike-by-spike (SbS) [3] [4]. Despite the differences in their underlying computational mechanisms we find that all three model classes achieved equivalent performance on a variety of tasks. This reveals that there are multiple –still unexplored– approaches to implementing deep learning algorithms.

Each of these model classes has its own advantages:

CNNs are the de-facto industry standard and are known to be very accurate at image recognition and classification.

NNMF [5] [6] is a technique used to decompose a non-negative matrix *X* as the product of two non-negative matrices *W* and *H*, such that *X ≈ WH*. The matrices *W* and *H* are chosen to have lower rank than *X*, which means that they have fewer rows or columns. This allows NNMF to extract a compact representation of the data in *X*, by identifying patterns and structures in the data. This decomposition can be used to extract useful features from the data represented by the original matrix. NNMF is often used for data analysis and dimensionality reduction, and has applications in fields such as image processing, text mining, and bioinformatics [8].

Deep networks based on non-negative matrix factorization (NNMF) have the advantage of producing factor matrices that are non-negative and have a clear interpretability as basis vectors for the latent factors.

Deep spiking convolutional neural networks (SNNs) [12] have several benefits over normal deep CNNs [9]. SNNs are designed to mimic the operations in brains which use sparse and asynchronous discrete signals for communication and processing. This allows them to operate in an event-driven manner, which can result in significant energy savings when implemented on neuromorphic hardware.

Converting spiking neural networks into event-based neuron models can also improve their ability to process spatiotemporal data. Event-based neuron models are designed to operate in an asynchronous manner, which allows them to process temporal data more effectively than traditional integrate-and-fire models.

“Spike-by-spike” (SbS) [3] is an event-based stochastic deep spiking neural network that allows for both training via back-propagation and unsupervised training of the weights. The latter means that the network can learn to adjust its weights based on the input data it receives, without the need for explicit supervision or labeled data.

SbS can be seen as a special case of non-negative matrix factorization (NNMF) where the continuous input distribution is replaced by individual stochastic spikes.

Stochasticity in deep neural networks can be helpful for increasing the robustness of classification by introducing noise into the network’s computations. This noise can help prevent overfitting by making it more difficult for the network to memorize the training data. Instead, the network must learn to extract generalizable features from the data that allow it to perform well on new examples. By introducing noise into the network’s computations, the network also becomes more robust to small perturbations in its inputs. This can be particularly useful for tasks that involve processing noisy or uncertain data.

## 2 Results

Both NNMF

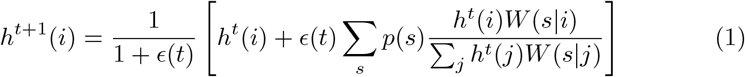

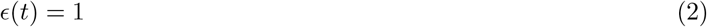

and SbS

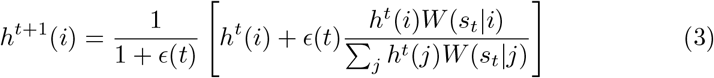

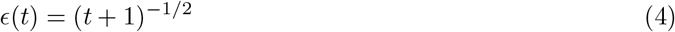

use an iterative dynamic on their latent variables *h*(*i*) to learn a representation of the data. This iterative process allows the network to adjust its internal representation of the data over time, based on the input it receives (the input distributions *p*(*s*) or the spikes *s*_*t*_ that are the indices of input neurons, which are generated stochastically from *p*(*s*)). For keeping the performances as comparable as possible we used the same network architecture for all three types of models (see figure 1). The stochastic spikes (i.e. neuron indices) are used for the *input spike layer* (see section 4.1) as well as for the communication between SbS layers.

**Fig. 1.**
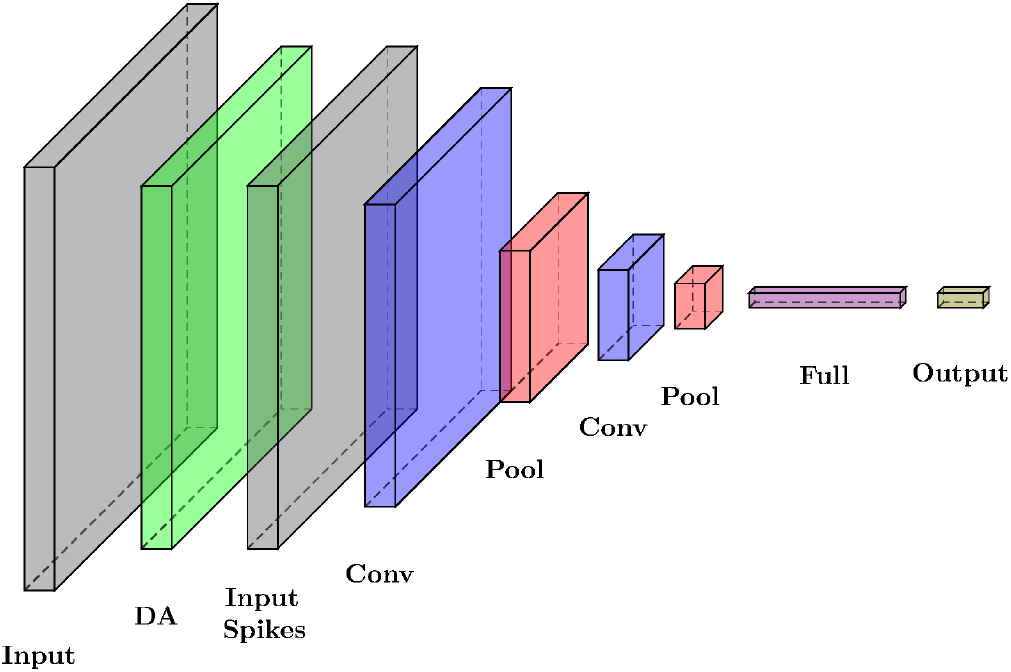
The network architecture. In the case of NNMF and SbS networks, the convolution layer (conv), the full layer, and the output layer are constructed from IPs with their according dynamic (equation 1 and 3) on their latent variables (see [4] for more details). For the CNN, the convolution layers, and the full layer are followed by ReLU layers. Also the pooling layers are max pooling layers instead of average pooing layers used for NNMF and SbS. The convolution layers, full layer and output layer have activated bias parameters. MNIST, n-MNIST, Fashion MNIST, and CIFAR10. 2D Convolution layer 1: kernel:5×5×32; stride:1×1 (valid). 2D Convolution layer 1: kernel:5×5×64; stride:1×1 (valid). Both 2D pooling layers: kernel:2×2; stride:2×2. CNNs use max pooling. NNMF and SbS uses average pooling. Fully connected layer: 96 neurons. Output 10 neurons. Input picture dimensions: *-MNIST: 28×28×1 and CIFAR10: 32×32×3.

For NNMF and SbS layers we introduced the so called inference populations (IPs) [4]. Every IPs operates independent of the other IPs in a layer. Inside an IP, the neurons are within a competition. While these networks could be trained via automatic differentiation, due to their iterative nature these would require excessive amounts of memory. We circumvented this by developing a special back-propagation learning rule which can be used for training deep NNMF and SbS networks. In section 4.4 we give a short recapitulation of the learning rule. More details can be found in [4]. For improving the speed of learning the weights, we added a scaling procedure for the back-propagated error (see section 4.3).

Further on, we observed that NNMF and SbS do not work well with the original Adam optimizer [1]. Therefore, we developed a modified version better suited for these type of networks (see section 4.5).

We tested all three network types with the so called On/Off-filtering [3]. This transformation is inspired by the on-off-cells of the visual system [7] and seems to improve the performance as well as noise robustness. It is used to split the input distribution *p*(*s*) into two channels *p*On(*s*) and *p*Off(*s*) using:

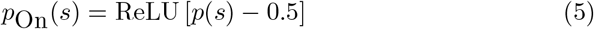

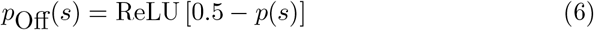

For the simulation we used the following datasets: MNIST, n(oise)-MNIST, Fashion MNIST, and CIFAR10. Information about the data-preprocessing and the data augmentation procedure are described in section.

Fig. 2 as well as the tables 1 and 2 allow to compare the results for the combinations of those datasets and the three network models with and without On/Off pre-processing.

**Fig. 2.**
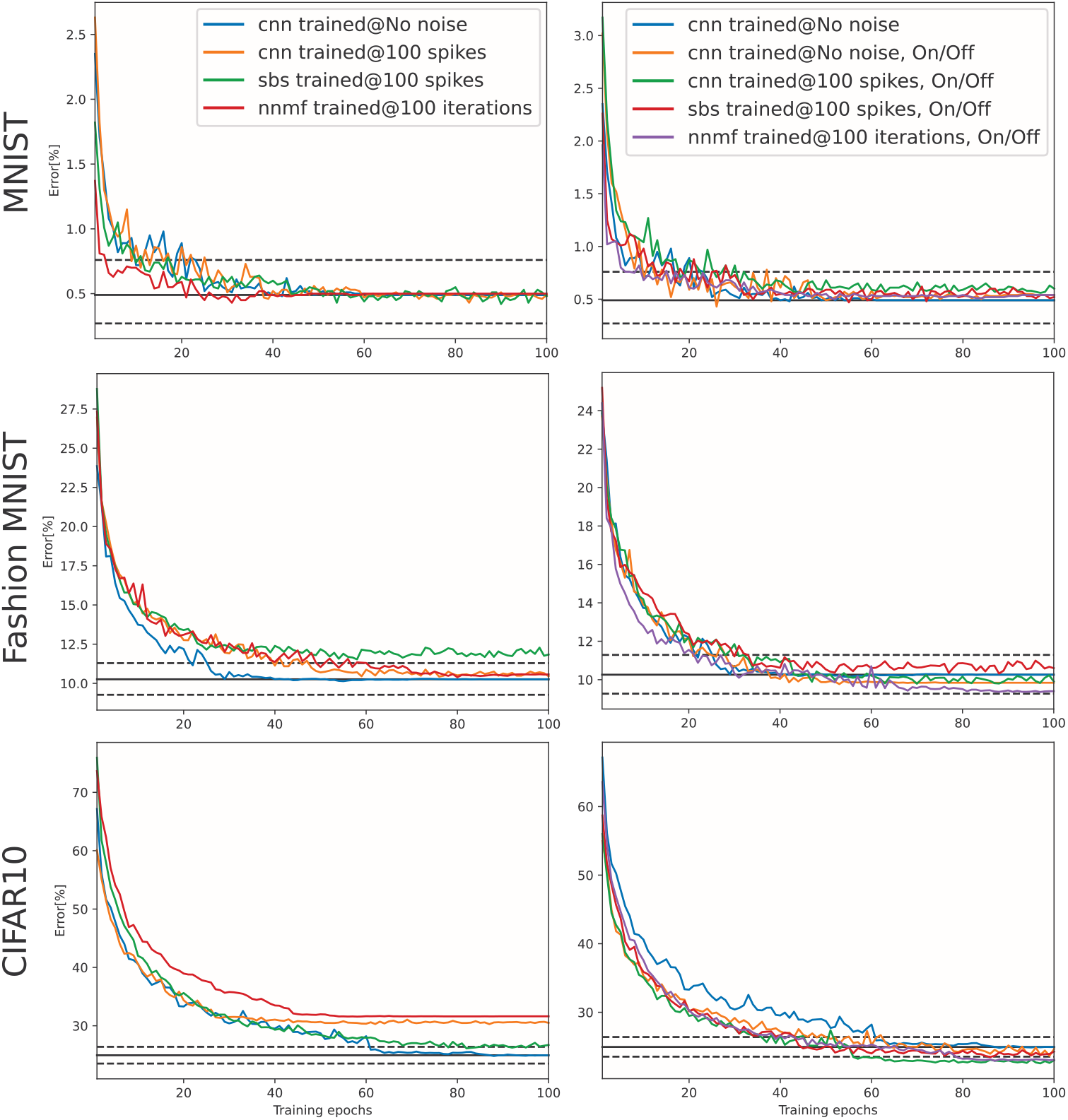
Convergence during training. Shown is the error on the test data. The solid black line is the final error (averaged over the last 10 epochs) of the CNN No Noise curve. See table 1 for the final performance values after training. The dashed black lines are the expected statistic deviations (p-value= 1*/*100) around this value, corresponding to the one sided Fisher exact test.

**Table 1.**
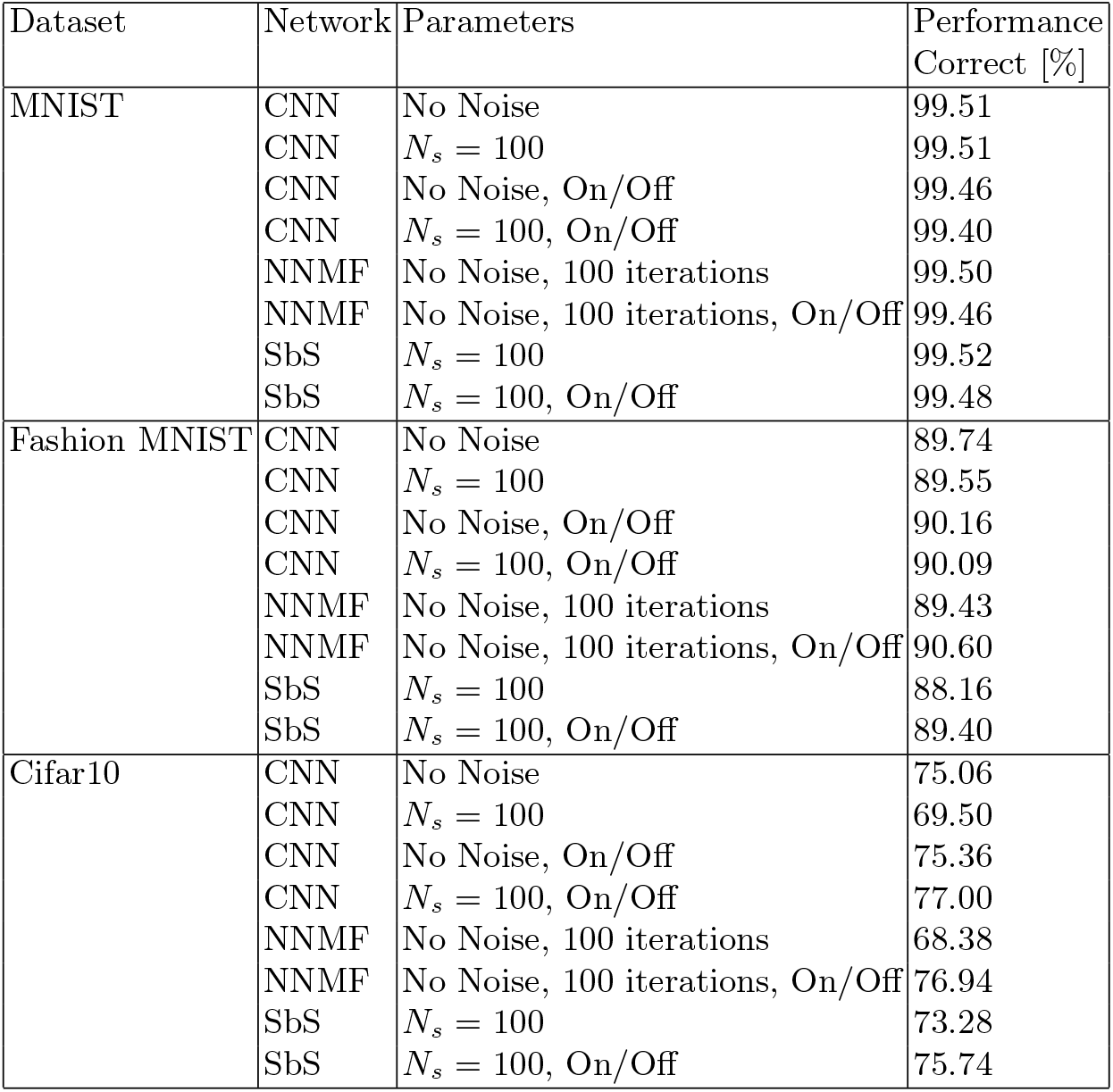
Final values of figure 2 (right column). The amount of spikes for the input layer generating spikes is *N*Spikes = *N*_*s*_ · *N*_*C*_ · *N*_*x*_ · *N*_*y*_. The On/Off split for the input neurons does not change *N*_*S*_.

| Dataset | Network | Parameters | Performance<br>Correct [%] |
| --- | --- | --- | --- |
| MNIST | CNN | No Noise | 99.51 |
| | CNN | $N_s = 100$ | 99.51 |
|  | CNN | No Noise, On/Off | 99.46 |
| | CNN | $N_s = 100$, On/Off | 99.40 |
|  | NNMF | No Noise, 100 iterations | 99.50 |
|  | NNMF | No Noise, 100 iterations, On/Off | 99.46 |
| | SbS | $N_s = 100$ | 99.52 |
| | SbS | $N_s = 100$, On/Off | 99.48 |
| Fashion MNIST | CNN | No Noise | 89.74 |
| | CNN | $N_s = 100$ | 89.55 |
|  | CNN | No Noise, On/Off | 90.16 |
| | CNN | $N_s = 100$, On/Off | 90.09 |
|  | NNMF | No Noise, 100 iterations | 89.43 |
|  | NNMF | No Noise, 100 iterations, On/Off | 90.60 |
| | SbS | $N_s = 100$ | 88.16 |
| | SbS | $N_s = 100$, On/Off | 89.40 |
| Cifar10 | CNN | No Noise | 75.06 |
| | CNN | $N_s = 100$ | 69.50 |
|  | CNN | No Noise, On/Off | 75.36 |
| | CNN | $N_s = 100$, On/Off | 77.00 |
|  | NNMF | No Noise, 100 iterations | 68.38 |
|  | NNMF | No Noise, 100 iterations, On/Off | 76.94 |
| | SbS | $N_s = 100$ | 73.28 |
| | SbS | $N_s = 100$, On/Off | 75.74 |

**Table 2.** Results for the n(oise)-MNIST dataset. Two variants are shown in this table: The first performance column represents the case were the networks were trained on the noisy data from scratch. The second performance column uses the already trained MNIST network shown in table 1 for inference.

| Dataset | Network | Parameters | Performance<br>Correct [%] | MNIST<br>weights<br>Correct [%] |
| --- | --- | --- | --- | --- |
| additive white<br>gaussian noise<br>(awgn) | CNN | Perfect | 98.87 | 48.18 |
| | CNN | $N_s = 100$ | 98.88 | 67.96 |
|  | CNN | No Noise, On/Off | 99.04 | 84.30 |
| | CNN | $N_s = 100$, On/Off | 99.10 | 90.75 |
|  | NNMF | No Noise, 100 iterations | 99.04 | 80.80 |
|  | NNMF | No Noise, 100 iterations, On/Off | 99.01 | 96.08 |
| | SbS | $N_s = 100$ | 98.98 | 89.18 |
| | SbS | $N_s = 100$, On/Off | 99.01 | 96.83 |
| motion-blur | CNN | No Noise | 99.42 | 96.57 |
| | CNN | $N_s = 100$ | 99.31 | 97.18 |
|  | CNN | No Noise, On/Off | 99.50 | 94.54 |
| | CNN | $N_s = 100$, On/Off | 99.38 | 94.11 |
|  | NNMF | No Noise, 100 iterations | 99.36 | 96.35 |
|  | NNMF | No Noise, 100 iterations, On/Off | 99.26 | 87.81 |
| | SbS | $N_s = 100$ | 99.37 | 96.79 |
| | SbS | $N_s = 100$, On/Off | 99.40 | 96.07 |
| reduced<br>contrast<br>and awgn | CNN | No Noise | 97.42 | 16.96 |
| | CNN | $N_s = 100$ | 97.53 | 36.12 |
|  | CNN | No Noise, On/Off | 97.69 | 79.25 |
| | CNN | $N_s = 100$, On/Off | 97.70 | 78.82 |
|  | NNMF | No Noise, 100 iterations | 97.89 | 64.99 |
|  | NNMF | No Noise, 100 iterations, On/Off | 98.04 | 79.32 |
| | SbS | $N_s = 100$ | 97.37 | 70.57 |
| | SbS | $N_s = 100$, On/Off | 97.87 | 85.18 |

With On/Off encoding of the inputs all models achieve equivalent performance (*p* = 0.01, see Fig. 2, table 1). For noisy MNIST (n-MNIST) table 2 reveals substantial noise robustness of the proposed approach: SbS-networks outperform both NNMF-networks as well as CNNs when trained on noiseless data and tested with corrupted inputs (compare the performance columns).

## 3 Discussion and Conclusion

SbS networks provide a trade-off between the non-spiking CNNs and networks based on integrate-and-fire neurons (IaF) [13]. CNNs run faster than SbS networks in current hardware architectures, but CNNs do not have the advantages associated with spikes [9]. IaF networks are biologically more realistic than SbS networks, but require orders of magnitude more computational power than SbS [10]. Furthermore, it is possible to construct very efficient hardware accelerators for SbS [11]. A Github with our PyTorch SbS and NNMF framework can be found at https://github.com/davrot/pytorch-sbs

We showed that the modified Adam optimizer combined with the error back-propagation rule and back-propagation scaling optimize performance for NNMF and SbS networks [4]. Also, we found that the on/off filtering improves the performance of NNMF and SbS networks (and sometimes even the performance of CNNs) which allows NNMF and SbS network to reach performance equivalent to CNNs. These findings hold for the three tested databases: MNIST, Fashion MNIST, and CIFAR10 (see table 1 and figure 2). Experiments with deeper neural networks and larger data sets have been denied to us by the limits of our available computing resources.

We used the n-MNIST dataset as test data for a network that was trained on the none noisy MNIST dataset. We found: a.) adding On/Off filtering to a CNN improves its noise robustness. b.) NNMF and SbS networks generalize better: they can handle different types of perturbations well even though they were never trained on this kind of corrupted data (see table 2, right column).

The fact that networks based on non-negative matrix factorisation can achieve the performance of standard CNNs proves that they are a valuable alternative to the usual deep CNNs. They have the advantage of interpretability, show superior noise robustness and, with their incremental inference of latent variables, could become fruitful for processing also temporal data. Their biologically realistic implementations in the event-driven spike-by-spike network SbS do not only achieve comparable performance levels, but also show superior noise robustness even when trained on noiseless data. Thus, SbS networks represent a major step towards interpretable neural networks, provide a promising framework for efficient hardware implementations and have the potential to also explain the surprising performances of real brains.

## 4 Methods

### 4.1 Input Spike layer

In SbS, the layer called *input spike layer* produces stochastic spikes based on the input distribution *p*(*X, Y, C*). A spike *s*_*t*_ is equivalent to the index of a neuron. The index of the neuron is determined by drawing randomly from the input distribution of a given input pattern after data augmentation. The number total of indices (spikes) generated by the input layer is defined as *N*Spikes, where *N*Spikes = *N*_*s*_ · *N*_*C*_ · *N*_*x*_ · *N*_*y*_. Here, *N*_*X*_ and *N*_*Y*_ represent the number of pixels in *X* and *Y* dimension, *N*_*C*_ represents the number of input channels (the on/off transformation does not double *N*_*C*_) and *N*_*S*_ represents the average number of spikes per input neuron. For CNNs, the generated spikes given the *X* and *Y* position as well as the channel *C* are counted and normalized into a new 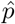(*X, Y, C*) distribution. In the case of SbS, spikes are send to the corresponding IPs according the convolution windows.

For *N*_*S*_ = 0 (i.e. No Noise), no spikes are drawn and *p*(*s*) is used directly. Also for all simulation with NNMF this is the case.

### 4.2 Data augmentation (DA)

Dimensions of the input pattern after data augmentation:

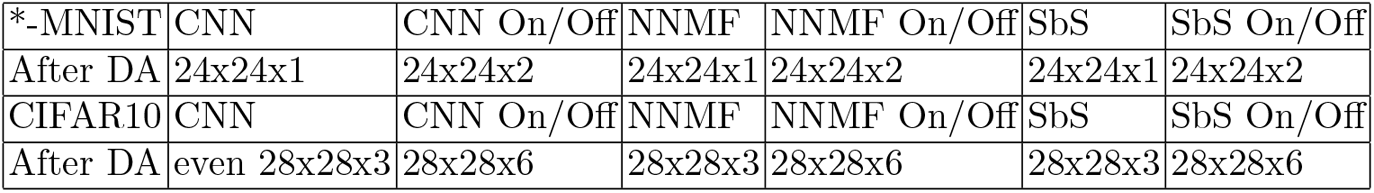

The employed data augmentation strategy for the different data sets was:

#### MNIST

http://yann.lecun.com/exdb/mnist/ and **n-MNIST** https://csc.lsu.edu/~saikat/n-mnist/

The gray-value images from the datasets MNIST and the noise-MNIST have the dimensions 28×28 pixel with 1 color channel. After loading the data, the images are converted into float32 values and normalized to the maximum value of the respective dataset. The PyTorch Vision procession chain was:

Training data:

|  |
| --- |
| Image |
| RandomCrop(size=(24,24)) |

Test data:

|  |
| --- |
| Image |
| CenterCrop(size=(24,24)) |

#### Fashion MNIST

https://github.com/zalandoresearch/fashion-mnist

The gray-value images have the dimension 28×28 pixel with 1 color channel. After loading the data, the images are converted into float32 values and normalized to the maximum value of the respective dataset. The PyTorch Vision procession chain was:

Training data:

|  |
| --- |
| Image |
| RandomCrop(size=(24,24)) |
| RandomHorizontalFlip(p=0.5) |
| ColorJitter(brightness=0.5, contrast=0.1, saturation=0.1, hue=0.15) |

Test data:

|  |
| --- |
| Image |
| CenterCrop(size=(24,24)) |

#### CIFAR 10

https://www.cs.toronto.edu/~kriz/cifar.html

After loading the data, the images are converted into float32 values and normalized to the maximum value of the dataset. The images have the dimensions 32×32 pixels with 3 color channels. In the case of the on/off filtering, every color channel is transformed individually and then concatenated into 6 color channels. The PyTorch Vision procession chain was:

Training data:

|  |
| --- |
| Image |
| RandomCrop(size=(28,28)) |
| RandomHorizontalFlip(p=0.5) |
| ColorJitter(brightness=0.5,contrast=0.1,saturation=0.1,hue=0.15) |

Test data:

|  |
| --- |
| Image |
| CenterCrop(size=(28,28)) |

### 4.3 Back-propagation error normalization

In the case of SbS and NNMF networks, the incoming back-propagation errors for each layer are normalized by dividing them by the maximum absolute value for that layer during the first mini-batch. Let 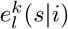 be the incoming error for layer *l* and mini-batch *k* and 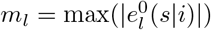 be the maximum absolute value for layer *l*. The normalized back-propagation errors for layer *l* can be written as 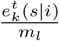. By normalizing the incoming errors in this way, the gradients 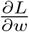 are kept within a reasonable range and issues with vanishing or exploding gradients can be avoided.

The training for all network types ran over 100 epochs, while one mini-batch contained 480 pattern each.

### 4.4 SbS and NNMF: backprop learning

#### Loss functions

For the CNN we used *torch*.*nn*.*functional*.*cross entropy* as *L*_*µ*_(*i*). For SbS and NNMF we used

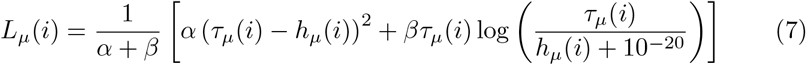

with *τ*_*µ*_(*i*) as the one hot target generate from the label. We used *α* = 0.5 and *β* = 1.

#### Backprop learning [4]

The backpropagation learning algorithm is described in detail in [4]. Here only a summary:

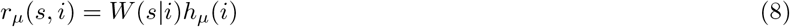

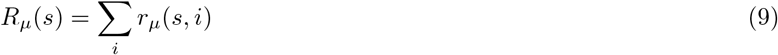

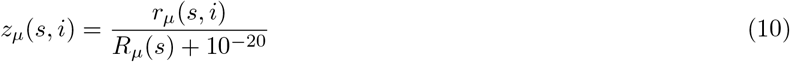

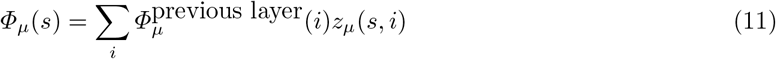

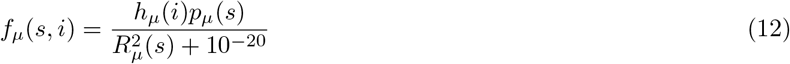

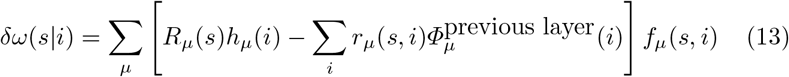

*p*_*µ*_(*s*) is the input into an IP. *µ* represents the pattern id, *x* and *y* coordinate.

#### Updating weights

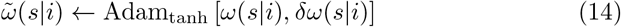

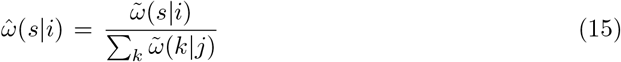

Find ever 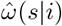(*s*|*i*) that is smaller than the threshold 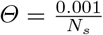 by *Θ*.

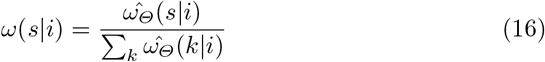

### 4.5 Adam optimizer

The Adam optimizer [1] is used to update the weights of neural networks during training. It is based on the concept of momentum and uses two moving averages to keep track of the first and second moments of the gradients. The first moment is an estimate of the mean of the gradients, while the second moment is an estimate of the uncentered variance of the gradients.

The Adam optimizer uses these moving averages to calculate an adaptive learning rate for each weight in the neural network. The adaptive learning rate is calculated by dividing the first moment by the square root of the second moment. This can help prevent oscillations and accelerates convergence.

The Adam optimizer calculates the weight updates as follows:

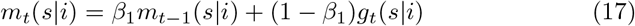

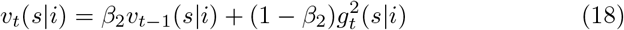

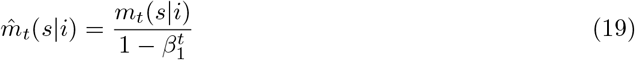

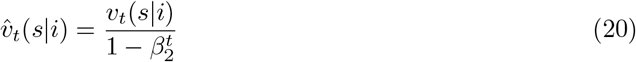

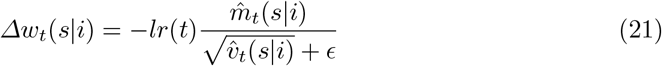

where *m*_*t*_ and *v*_*t*_ are estimates of the first and second moments of the gradients, 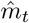 and 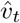 are bias-corrected estimates of the first and second moments, *lr*(*t*) is the learning rate at time *t, β*_1_ = 0.9 and *β*_2_ = 0.999 are exponential decay rates for the first and second moment estimates, and *ϵ* = 10^*−*8^ is a small constant to prevent division by zero.

For the SbS and NNMF network training we modified this method. Instead of directly using the weight updates calculated by the Adam optimizer, we first apply the tanh function to the weight updates. By applying the tanh function to the weight updates, we can limit the range of weight updates to -1 to 1. Next, we scale and shift the output of the tanh function by multiplying it by 0.5 and adding 1.0. This changes the range of the weight updates to 0.5 to 1.5. This ensures that the weight updates consist of only positive numbers. If the initial weight matrix has only positive values, then after applying this method, the updated weights will still be positive. This ensures that the non-negativity constraint on the weights is satisfied.

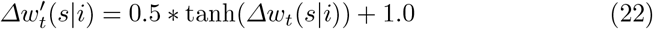

The updated weights are then calculated as follows:

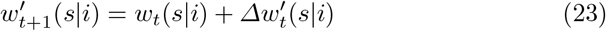

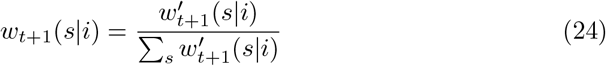

### 4.6 Learning rate scheduler

To control the learning rate *lr*(*t*) the PyTorch learning rate scheduler *ReduceL-ROnPlateau* was used. This scheduler reduces the learning rate when the monitored loss *L*(*t*) has stopped improving. The initial learning rate is *lr*(0) = 0.001 for CNNs and *lr*(0) = 0.01 for SbS and NNMF networks. If the scheduler sees no improvement for *L*(*t*) over 10 epochs the learning rate is reduced by a factor of 10.

## 5 Acknowledgement

We acknowledge support by the following grants: DFG: Efficient implementation of spike-by-spike neural networks using stochastic and approximate techniques (PA 569/6-1, GA 763/15-1), DFG SPP: Evolutionary optimisation of neural systems (SPP 2205) https://gepris.dfg.de/gepris/projekt/402741184 – Evolution of flexibility - optimisation of task-dependent information processing in the visual system (ER 324/5-1), Era-Net Neuron https://www.neuron-eranet.eu: I-See – Improved intra-cortical visual prostheses through complex coding and integration of spontaneous activity states (BMBF 01EW2104A), Stiftung Bremer Wert-papierboerse https://www.stiftung-bwb.de.

During writing we took advantage from special tools including Bing Chat https://www.bing.com and DeepL https://www.deepl.com.

